# FungAMR: A comprehensive portrait of antimicrobial resistance mutations in fungi

**DOI:** 10.1101/2024.10.07.617009

**Authors:** Camille Bédard, Alicia Pageau, Anna Fijarczyk, David Mendoza-Salido, Alejandro J. Alcañiz, Philippe C. Després, Romain Durand, Samuel Plante, Emilie M. M. Alexander, François D. Rouleau, David F. Jordan, Adarsh Jay, Mathieu Giguère, Mégane Bernier, Jehoshua Sharma, Laetitia Maroc, Nicholas C. Gervais, Anagha C.T. Menon, Isabelle Gagnon-Arsenault, Sibbe Bakker, Johanna Rhodes, Philippe J. Dufresne, Amrita Bharat, Adnane Sellam, Domenica G. De Luca, Aleeza Gerstein, Rebecca S. Shapiro, Narciso M. Quijada, Christian R. Landry

## Abstract

Antimicrobial resistance (AMR) is a global threat. To optimize the use of our antifungal arsenal, we need rapid detection and monitoring tools that rely on high-quality AMR mutation data. Here, we performed a thorough manual curation of published AMR mutations in fungal pathogens to produce the FungAMR reference dataset. A total of 501 papers were curated, leading to 35,792 mutation entries all classified with the degree of evidence that supports their role in resistance. FungAMR covers 95 species, 246 genes and 208 drugs. We combined variant effect predictors with FungAMR resistance mutations and showed that these tools could be used to help predict the potential impact of mutations on AMR. Additionally, a comparative analysis among species revealed a high level of convergence in the molecular basis of resistance, highlighting some potentially universal resistance mutations. The analysis also showed that a significant number of resistance mutations lead to cross-resistance within antifungals of a class, as well as between classes for certain mutated genes. The acquisition of fungal resistance in the clinic and the field is an urging concern. Finally, we provide a computational tool, ChroQueTas, that leverages FungAMR to screen fungal genomes for AMR mutations. These resources are anticipated to have great utility for researchers in the fight against antifungal resistance.

## Introduction

Infections caused by fungal pathogens constitute a significant health threat and socio-economic burden. Fungal pathogens have the potential to cause severe diseases, accounting for multiple million deaths annually^1^. The mortality rate of patients with invasive mycosis, an infection spreading within organs, can exceed 50%^2^. Yet, fungal infections are amongst the most overlooked infectious diseases. The World Health Organization (WHO) estimated that less than 1.5% of the funding allocated for infectious disease research is directed towards investigating fungal infections^3^. The burden of pathogenic fungi extends beyond human infections, encompassing challenges to both food security and biodiversity. Fungi are a leading cause of plant diseases, resulting in an estimated 20% worldwide crop yield loss^4^. Additionally, certain fungi have been responsible for significant declines in animal populations, such as amphibian and bat populations across the globe^5,6^, and few if any effective drugs are available to mitigate critical loss of animal biodiversity.

Another pressing threat to public health is antimicrobial resistance (AMR)^7^. The widespread use of antifungals in agriculture and clinics, or in other sectors as preservatives, has led to the emergence of resistant strains and populations unresponsive to treatments^8,9^. Fungicide resistance in plant pathogens exacerbates the threat to food security by increasing yield losses and contamination of crops with mycotoxins^9^. AMR in human fungal pathogens poses a particularly daunting challenge due to the limited treatment options available^8,10^. The medical community relies on only a handful major classes of antifungals to treat human systemic infections^11^. Even if other compounds are under development, treatment options are currently rapidly depleted when resistance mutations arise in patients during treatment^8,10^. This is exacerbated by the overlap between the classes of antifungals used to treat infections in humans and the fungicides used on plants, which selects for resistance mutations in environmental fungi that can go on to infect humans^4^. For instance, exposure of *Aspergillus fumigatus* to agricultural azole fungicides leads to cross-resistance to medical azole antifungals^12,13^.

One approach to overcome AMR in fungi could be to develop new drugs. However, humans and fungi share a much more recent common ancestor compared to humans and bacteria. This considerable proportion of shared genes and cell morphology limits the number of unique molecular targets that only affect the fungal pathogen and not the human host cells^14^. Despite ongoing efforts to extend our current antifungal arsenal and identify new targets, the rate at which new antimicrobials are produced is far exceeded by the speed at which resistance emerges^14^. Additionally, resistance will eventually likely emerge against these new drugs. While developing and discovering new antifungals is crucial, this alone will not address the problem of antifungal resistance. A better understanding of the different resistance mechanisms is essential in our battle against fungal pathogens to help guide the development of treatment strategies, and hopefully, to better understand and control the evolution of resistance. Having a centralized antifungal mutation repository of high quality will also improve our capacity to detect resistance and paves the way for routine genotypic detection of antifungal resistance (vs long and cumbersome phenotypic assays such as broth microdilution).

Unlike antibiotic resistance, which frequently emerges from the acquisition of genes, antifungal resistance emerges through *de novo* mutations in the genome resulting in a wide array of possible resistance mechanisms^15^. Accordingly, various mechanisms of AMR have been described for all available antifungal classes. Yet, the number of fungal AMR mutations documented in the literature remains relatively limited and dispersed, and in many cases we lack an understanding of the precise impact or causal nature of antifungal resistance mutations. In-depth knowledge of resistance mutations across all available drugs would allow us to better understand which mutations are directly linked with antifungal drug resistance, and further determine the extent of cross-resistance within and between antifungal classes. Similarly, knowledge of resistance mutations across different species and genes would allow us to assess how resistance mechanisms are conserved among fungi. Also, a better understanding of resistance mutations would enable us to streamline the development of antifungal agents that overcome resistance mutations and avoid cross-resistance characteristics by modifying currently available drugs. Developing new antifungals using this strategy would alleviate the pressure to discover new target genes. This knowledge could facilitate the development of genomic tools to monitor and interpret genetic variants of pathogens in real-time during treatment or in the agricultural field to detect and follow resistance. Such information could also be used to build predictive models to foresee resistance mutations before their spread in clinics and agriculture. Databases on antibacterial resistance mutations, such as CARD^16,17^, have proved invaluable in addressing the serious threat posed by AMR for instance, by enabling the interpretation of genomic information. However, while some antifungal resistance databases exist, these are unfortunately outdated^18^ or contain a large fraction of inaccurate information when done using simple text mining (Supplementary note 1)^19^. Since many studies are reporting potentially spurious associations between mutations and resistance, a careful annotation is needed to build a useful reference dataset (see below). Thus, there is a need to exhaustively and reliably gather all antifungal resistance mutations reported in the literature in one accessible repository.

To address these limitations, we manually curated 501 papers reporting fungal AMR mutations to create the FungAMR compendium (Supplementary table 1, <u>FungAMR Mutation Data</u>). FungAMR contains 35,792 carefully curated entries across 208 drugs (including 118 antifungals) for 95 fungal species and contains missense mutations as well as other genomic changes such as copy-number variations and nonsense mutations. Every mutation is classified with the degree of evidence that supports its role in AMR. We have taken advantage of this new resource to better understand resistance mechanisms among species and antifungals. We have combined variant effect predictions with resistance mutations to confirm that many genes confer resistance through loss of function mutations and show that such analyses could be used to help interpret the impact of mutations on AMR. Furthermore, comparative analysis of resistance mutations among species revealed a high level of convergence in resistance mechanisms at both the gene and the mutation levels. This analysis also confirmed that many resistance mutations provide cross-resistance to antifungals within the same class, but also between classes for mutations present in certain genes. Finally, we present *Chromosome Query Targets* (ChroQueTas), a computational pipeline that utilizes the information contained in FungAMR to easily detect the presence of AMR mutations in fungal genomes and produce an annotated report. Overall, the analysis of the content of FungAMR revealed biases in the study of certain species, proteins and antifungals, and highlighted where more research is needed. This resource will help the scientific community to address the serious threat of antifungal resistance, and has the potential to provide a launchpad for fundamental and applied research.

## Methods

### Literature review, data curation and development of the FungAMR resource

All papers were read by a scientist in the field and the information was extracted in a systematic manner. In addition to the laboratories involved, some of the curators were recruited among graduate students and postdoctoral fellows from laboratories working on fungal genetics and antifungal resistance, which are part of a pan-Canadian training program on the evolution of fungal pathogens, <u>EvoFunPath</u>. Two undergraduate students were involved in curating papers that more senior scientists also curated. Curation was conducted between April 2023 and March 2025 and included publications from 1988 to 2024. All curators collected papers reporting antimicrobial resistance using standard literature searching and mining references from other papers. Curators initially used general terms related to fungal antimicrobial resistance such as Resistance, Antifungal, Fungicide, Mutation, Candida, Aspergillus, Cryptococcus, Zymoseptoria, Azoles, Echinocandins, Polyenes, 5-fluorocytosine, Flucytosine, Fungi, SNP, Deletion, Overexpression, Aneuploidy, etc. Then, other relevant references were found in the research papers or reviews and included in the curation. We examined human, animal and plant pathogens in addition to model species such as the laboratory model *Saccharomyces cerevisiae*, which is used as a model system to study drug resistance. We also re-curated all papers reported in the MARDy database^18^ as these did not come with any measure of confidence levels (see below).

Each paper was read by one of the curators, and mutations reported to be associated with resistance were extracted. Mutations also reported to likely not cause resistance were included in FungAMR, but not used in the analysis since it is difficult to judge whether this is useful information as assays rarely contain positive controls in parallel. Each entry (Supplementary table 1) is associated with the first author, the journal, the publication year and the PubMedID (PMID) of the paper in which the mutation was reported, the fungal species and its host, the gene name and its protein’s accession number, the location of the mutation, its impact on the gene (deletion, insertion, amino acid change, duplication) and the impact on susceptibility to the drugs assayed. We also recorded the origin of the strains (environmental, clinical, laboratory or evolved) as well as the quantitative measures of resistance or susceptibility when provided by the authors. These were often minimum inhibitory concentrations (MIC), but many more measures were reported such as the half maximal effective concentration (EC50), a more common quantitative measure used in the agricultural area. We tried to record these measures as faithfully as possible. Still, we found that the diversity of methods used made any systematic recording of the quantitative impact of the mutations difficult. We therefore relied on the author’s assessment to record the mutation as being associated, or not, with resistance. When the authors did not assess if a mutation was associated with resistance, we define strains as resistant when they have a significantly higher growth in the presence of an antifungal than the reference strain or when they present with a minimal inhibitory concentration to an antifungal above the established breakpoint. Each reported entry concerned a particular strain and its associated phenotype. For instance, if a strain was reported and had multiple amino acid substitutions in the gene or genes sequenced, those multiple substitutions were reported together in a single entry as it was impossible to discern which one could have been causal. When available, we also recorded the identifier of the genes reported so we could track the reference sequences for species of interest. These identifiers came from various databases such as the Candida Genome Database or NCBI. However, this was not systematically done by the authors so some entries are left blank. Not all papers reported the gene identifier for the reference genes the mutations were reported to impact. We therefore performed a quality check by comparing the mutations reported (change from WT genotype to mutant one) with the standard reference genes from the species of interest (Supplementary table 2). A commentary column is included where curators noted information relevant to the entry, such as the experimental method, the culture media or the antifungal susceptibility testing method (e.g EUCAST, CLSI). Finally, we extracted from all curated papers any information relevant to the clinical outcome, this information is listed in the “clinical outcome” column.

All curators annotation sheets were combined into a single table, after refining and editing the nomenclature of species, genes or proteins, mutations, drugs, journal names, and strain origins, to facilitate the comparison of mutations. MIC values were edited and converted to the same units whenever possible. Two additional columns were added, one with an alternative species name and another one with a gene name that corresponds to orthologs or homologs of that gene in other species. In case when the study was curated by multiple curators, entries of both curators were first compared for consistency and corrected if necessary. Redundant entries from the same study (mutations in the same gene, drug, and strain) were combined or one of the duplicate entries was removed.

For each mutation or group of mutations, we assigned a score that assesses the degree of evidence that supports their role in drug resistance. The scoring scheme was inspired by the use of Multiplexed Assays of Variant Effects (MAVEs) to classify human pathogenic variants^20,21^ and based on examples of resistance reports that we had seen in the literature and were agreed upon and discussed among the curators. The list of confidence scores associated with their description is presented in Table 1. A positive confidence score denotes mutations reported to confer resistance, while a negative confidence score relates to mutations reported in susceptible strains. A low positive confidence score indicates that the evidence for the contribution of the mutation to resistance is strong, with 1 being the strongest and 8 being the weakest. Similarly for negative scores, the evidence of susceptibility is stronger for −1 than −8.

**Table 1.**
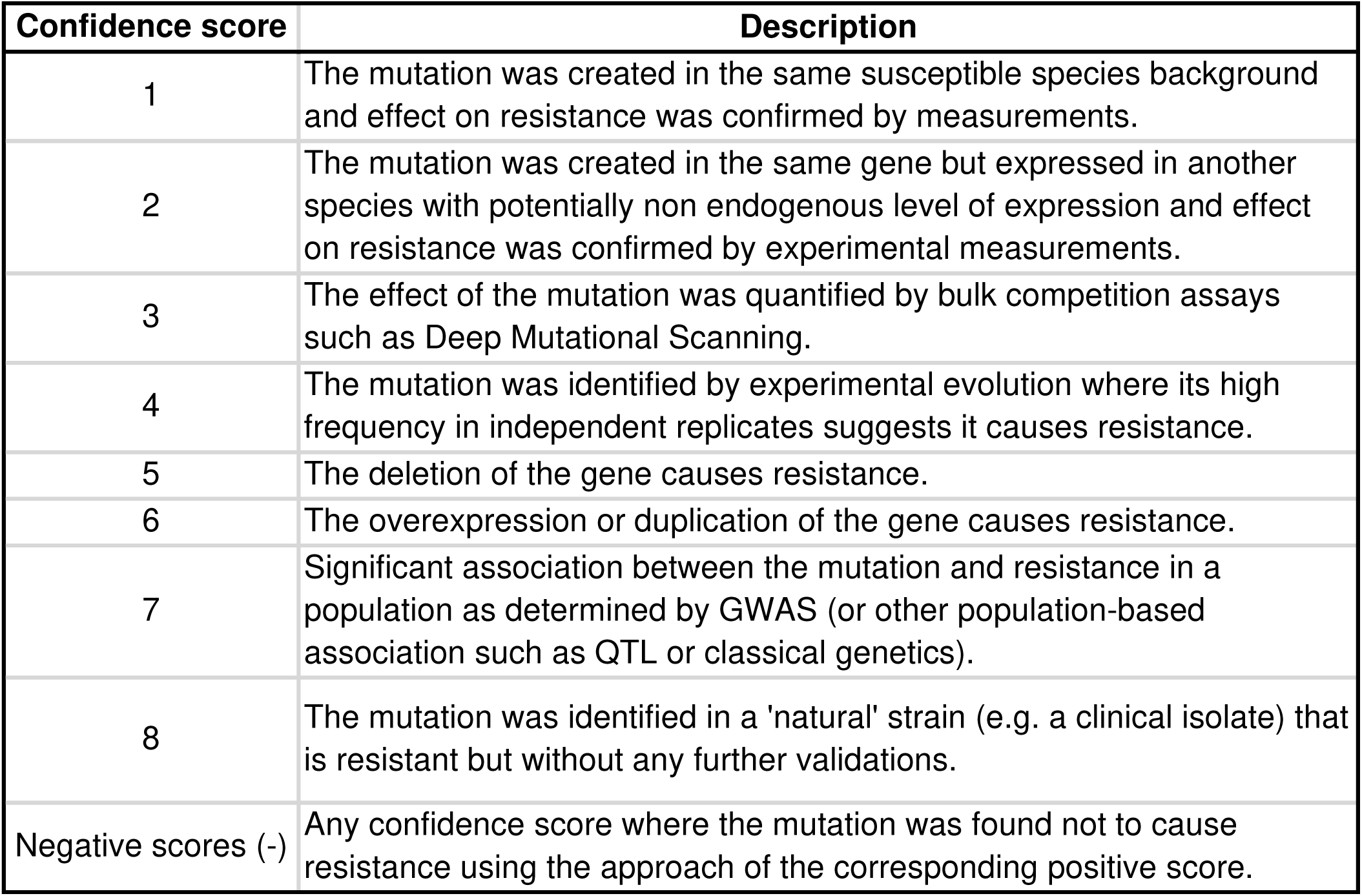
Description of the confidence scores associated with curated mutations in FungAMR.

| Confidence score | Description |
| --- | --- |
| 1 | The mutation was created in the same susceptible species background and effect on resistance was confirmed by measurements. |
| 2 | The mutation was created in the same gene but expressed in another species with potentially non endogenous level of expression and effect on resistance was confirmed by experimental measurements. |
| 3 | The effect of the mutation was quantified by bulk competition assays such as Deep Mutational Scanning. |
| 4 | The mutation was identified by experimental evolution where its high frequency in independent replicates suggests it causes resistance. |
| 5 | The deletion of the gene causes resistance. |
| 6 | The overexpression or duplication of the gene causes resistance. |
| 7 | Significant association between the mutation and resistance in a population as determined by GWAS (or other population-based association such as QTL or classical genetics). |
| 8 | The mutation was identified in a 'natural' strain (e.g. a clinical isolate) that is resistant but without any further validations. |
| Negative scores (-) | Any confidence score where the mutation was found not to cause resistance using the approach of the corresponding positive score. |

When a mutation is reported in several independent studies or strains within a study, the strongest resistance evidence (i.e. the best positive confidence score reported) and the strongest sensitivity evidence (i.e. the strongest negative confidence score reported) are provided in two separate columns.

### Variant effect predictors

For all protein reference sequences retrieved, we used HHblits^22^ from HH-suite3^23^ with each sequence as a query to create multiple sequence alignment (MSA) representative of the orthologs diversity. Initial HHblits iterative sequence search filter criteria were a minimum coverage of 80% with query sequence, at least 20% identity to the query and maximum 98% and the e-value cutoff for inclusion was 0.0001. Filter criteria were then gradually relaxed to get a minimum of 200 sequences in the final alignment. 267 alignments were produced with a median of 847 sequences per alignments. With these MSA files as input, we ran GEMME^24^ locally from docker desktop to compute the predicted effect of every single amino acid substitution across the proteins.

The protein structures from the proteins contained in FungAMR were obtained from the AlphaFold database (AlphaFold Protein Structure Database, Supplementary table 3). We used these structures to estimate the effects of amino acid substitutions on protein stability using FoldX^25^. The RepairPDB function was run 10 times with water prediction to repair incorrect torsion angles, VanderWaals clashes and total energy of the structure. We then used the MutateX Python package^26^ to perform an in-silico deep mutational scanning of the whole protein running the BuildModel function from FoldX and computing the difference in Gibbs free energy between the mutant and the wild-type.

### Tertiary protein structure predictions

When experimentally determined protein structures were available, they were retrieved from the Protein Data Bank (<u>RCSB PDB</u>, PDB). No complete tertiary protein structures were available on the PDB for Erg3, Hmg1, Upc2 and Pdr1, therefore we used the Alphafold 3^27^ web server (<u>AlphaFold Server</u>) to predict the entire structures (seed; Erg3: 382113193, Hmg1: 1841159774, Upc2: 601343710 and Pdr1: 264428788). Protein sequences were retrieved from UniProt (<u>UniProt</u>, Erg3: P32353, Hmg1: Q4WHZ1, Upc2: Q59QC7 and Pdr1: P12383). For Hmg1, the NAP – nicotinamide-adenine-dinucleotide-phosphate ligand was included in the prediction. For the transcription factors Upc2 and Pdr1, a DNA sequence containing one of the described DNA binding site of the enzymes was included in the prediction (Upc2: AATATCGTACCCGATTATGTCGTATATT from *C*. *albicans ERG11* promoter^28,29^ and Pdr1: GAGAAATGTCTCCGCGGAACTCTTCTAC from *S*. *cerevisiae PDR5* promoter^30^).

### Overall data description

We constructed tables of correspondence of amino acid sequences among the orthologs of the proteins reported in the data. Reference sequences for all proteins were retrieved from the Uniprot or NCBI database and MSA were performed on orthologous sequences using Muscle5^31^. All of the alignments for each family of orthologs are available in Fasta format as supplementary material (Supplementary data 1). These MSA files were used to assign a group number to orthologous mutations in different species, which can be found in an additional column. Alignment positions were also used to produce figures and to map resistance mutations on protein structures. The classification of the host of the fungal species, drug classes and gene types used in the figures is available in Supplementary table 4.

To display phylogenetic relationships among the species present in FungAMR, we used a phylogenetic tree constructed by Li et al.^32^ which includes 67 species from the database. To include approximate positions of the 18 remaining taxa at the species level, subphylogenies of species subsets were built using ITS sequences from either ITS RefSeq (BioProject number PRJNA177353), UNITE database v9, or if these were missing, from GenBank. Specific host forms (forma specialis) and hybrids were omitted from the species tree. Any ambiguous bases were resolved randomly, then sequences were aligned with mafft v7.525^33^ and trimmed using trimAl v1.4^34^ with “-gappyout” option. The maximum likelihood trees were built with iq-tree v2.2.2.7^35^. Branches with missing species were added to the main phylogenetic tree retaining the proportion of the branch lengths and position of the nodes with respect to related species from the ITS tree. The visualization of the tree was made using ggtree package^36^ in R.

The visualization of protein structures was done with ChimeraX^37^. Figures and analyses were done using the following Python packages: Matplotlib^38^, NumPy^39^, SciPy^40^, pandas^41^ and seaborn^42^, and the R packages: dplyr^43^, tidyr^44^, stringr^45^, igraph^46,47^, ape^48^, castor^49^, RRphylo^50^, and ggplot2^51^.

### Computational pipeline to detect mutations and InDels causing resistance

In addition to the FungAMR repository, a bioinformatic tool named *Chromosome Query Targets* (ChroQueTas) was developed in order to rapidly screen for AMR mutations in fungal genomes. ChroQueTas was devised to work CLI-based on UNIX environments (tested on Debian and Ubuntu-based OS) in a user-friendly manner so it can be run by following straightforward instructions and commands. ChroQueTas is open-source and publicly available at https://github.com/nmquijada/ChroQueTas, together with instructions for its installation and usages.

ChroQueTas requires an assembled fungal genome (either complete or draft assembly in FASTA format, no need for annotation) and a “scheme” (to be provided by the user under the -- *scheme* flag) corresponding to the species of the fungal genome (thus corresponding to the species reported in the FungAMR repository, and available by using the *--list_schemes* flag). With that information, ChroQueTas will: i) extract from the fungal genome the CDS and protein where a point mutation is known to cause AMR in that particular species by using miniprot v0.14-r265^52^ and the information contained in FungAMR (the translation genetic code is automatically set to “standard” for most fungi and to the “alternative yeast code” for the CTG clade, but can be customized by enabling the *--trans_code* flag; ii) evaluate sequence similarity against the reference by using BLASTP v2.14.1+ ^53^ and discard low confidence hits (to be specified by the user and the *--min-id*, *--min-cov* flags); iii) deal with potential introns, exons and InDels; iv) evaluate amino acid positions between the query and the reference proteins accounting for FungAMR information; and v) report amino acid changes and InDels that could lead to AMR according to the confidence score in FungAMR, together with their level of evidence and the predicted resistance phenotype. The output from ChroQueTas will consist of different text files with the information resulting from the AMR screening and further metadata contained in FungAMR, which is automatically downloaded and formatted by ChroQueTas during installation. ChroQueTas is intended to be a “living project”, being hosted in a public repository with enabled discussion forums where the users could ask for further improvements that will be considered and implemented on a regular basis.

In order to illustrate the power of the software, the Illumina raw sequencing data from 46 *Candida albicans* and 144 *Zymoseptoria tritici* isolates was downloaded using SRA Toolkit v3.1.0 (https://github.com/ncbi/sra-tools) (Supplementary Table 5). The sequencing data quality was assessed using FastQC v0.11.9^54^. Residual adapters and barcodes were discarded and quality filtering was performed by using FastP v0.23.2^55^ by setting a minimum Phred score of 25 and a minimum length of at least 75% of the length of the raw FASTQ files. Genomes were assembled by using SPAdes v3.15.5^56^ and contigs below 1,000 bp were discarded. The quality and completeness of each draft genome was assessed with QUAST v5.0.2^57^, BUSCO v5.4.3^58^ and by aligning the high-QC reads against the draft genomes by using Bowtie2 v2.4.2^59^. Once the quality of the genomes was assessed, ChroQueTas was used for the inference of AMR by using the following command:

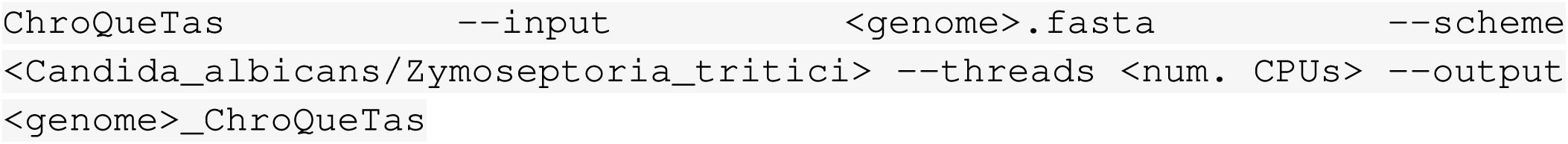

The average nucleotide identity (ANI) of the fungal genomes was calculated using fastANI v1.32^60^ and the distance matrix was imported into R environment^61^ to build a phylogenetic tree by using *ggplot2*^51^, *ggtree*^36^ and *reshape*^62^ packages.

## Results

### Content of FungAMR

A total of 501 papers were curated, leading to 35,792 mutation entries (Supplementary table 1, <u>FungAMR Mutation Data</u>). All entries are associated with information on the publication from which it was curated, as well as the gene, the location of the mutation, and the impact on drug susceptibility as experimentally assessed in the laboratory. Since the definition of a resistance mutation varies between studies, we defined a resistant strain as a strain presenting with a minimal inhibitory concentration to an antifungal above the established breakpoint when it was available or as a strain with significantly higher growth in the presence of an antifungal than the reference strain. Also, the level of evidence for the causal link between reported mutations and drug resistance varies considerably from one study to another. Some reports demonstrate a causal link between the mutation and the phenotype, for instance, by creating the mutation in a control genetic background. Other reports simply identify a mutation in a resistant strain, without further analysis, and often focus on sequencing only known resistance genes. Therefore, we associated all mutations or groups of mutations with a confidence score assessing the degree of evidence that supports their role in drug resistance (Table 1). A positive confidence score denotes mutations reported to confer resistance, while a negative confidence score relates to reported mutations that were not found to confer resistance. A low positive confidence score indicates that the evidence for the contribution of the mutation to resistance is strong, with 1 being the strongest and 8 being the weakest. We do not use the negative score in our analysis as those may depend on many factors, including the lack of sensitivity of the assays. Nevertheless, we think this information is useful to have recorded. When a mutation is associated with both positive and negative scores, we take the lowest positive one for our analyses. Ultimately, it is important to bear in mind that confidence scores do not translate into a probability of treatment failure, but only reflect the level of evidence we have that a mutation confers resistance in laboratory conditions. When available in a study, we listed any information relevant to the clinical outcome in the “clinical outcome” column.

We gathered more than 10,000 unique mutations for 208 drugs across 246 genes and 95 fungal species. Mutations assayed by Deep Mutational Scanning (DMS, confidence score of 3) come from studies where complete or nearly complete libraries of gene mutants were subjected to resistance testing. We have only four studies reporting mutations assayed by DMS, but they represent 76% of the entries (3: 21% and −3: 55%, Fig. 1a), showing how powerful systematic approaches can be at characterizing mutations. We set aside these entries for some analyses, since they are not representative of the overall content. Although DMS studies contribute to a large fraction of the data, 498 studies report non-DMS data for 8,568 entries and over 3,000 unique mutations covering the same set of species. The drugs listed include 42 antifungals used in the clinic (81% of entries), 76 used in agriculture (13% of entries) and 86 other molecules (6% of entries, e.g. anticancer drugs and immunosuppressants). However, most reported entries are of low confidence, with 58% having a confidence score of 8 (8: 43% and −8: 14%, Fig. 1a). Around 20% of the entries are mutations validated directly in the susceptible pathogen (1: 11% and −1: 4%, Fig. 1a) or by heterologous expression in the model yeast *Saccharomyces cerevisiae* (2: 4% and −2: 1%, Fig. 1a). This highlights the fact that although many AMR mutations have been reported in the literature, most have very little to no experimental support.

**Fig. 1.**
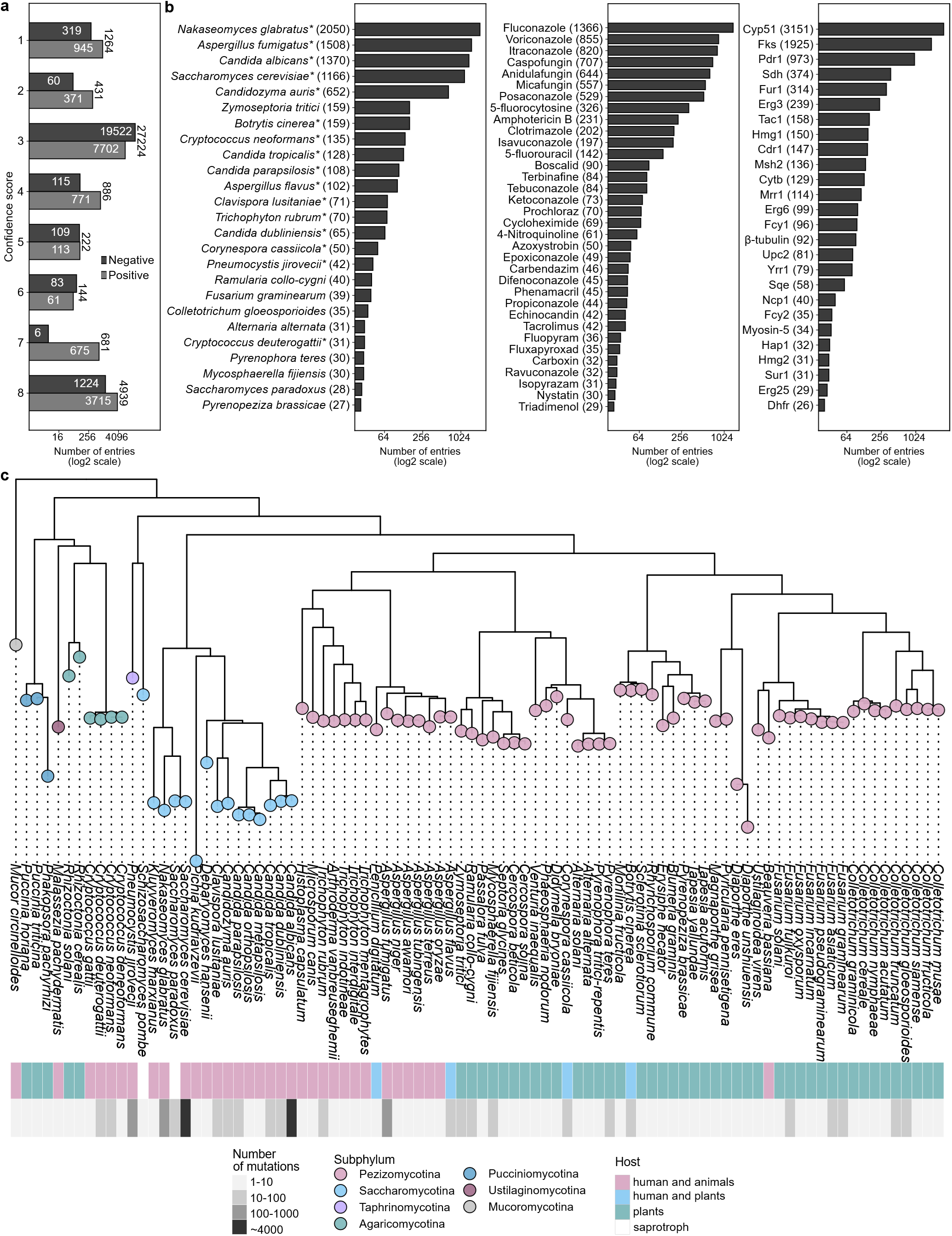
Content of FungAMR per confidence scores, species, antifungals and proteins. **a,** Distribution of positive and negative confidence scores associated with mutations reported in the literature (*n* = 54, 666 entries). A positive confidence score indicates mutations that have been reported to confer resistance, while a negative confidence score indicates mutations that have been reported but have not been found to confer resistance. A low positive confidence score means that the evidence for the effect of the mutation on resistance is strong, with 1 being the strongest and 8 being the weakest. Most entries come from high throughput experiments and have a confidence score of 3. Most remaining reports have limited experimental support. **b,** Number of observations per species, per drugs and per proteins. Species with * can cause human diseases (e.g. asthma, dermatosis, invasive mycosis) and drugs with * are used in clinics. Entries with a confidence score of 3 were excluded and cases with fewer than 25 reports are not shown to simplify the figure. **c,** Phylogenetic relationship of the species represented. On the right of species names are indicated which hosts these species can be isolated from and the number of unique mutations reported in the dataset in each case. The number of unique mutations can be lower than the number of mutation entries since many mutations have been reported multiple times.

*Nakaseomyces glabratus* (formerly *Candida glabrata*), *Aspergillus fumigatus and Candida albicans* have the largest number of entries with more than 1,300 each. They are followed by the model yeast *Saccharomyces cerevisiae* (1,166 entries) and the emergent human fungal pathogen *Candidozyma auris* (formerly *Candida auris*, 652 entries) (Fig. 1b). Roughly 75% of the listed species have less than 25 entries, reflecting a strong bias towards the study of certain fungi. Antifungals used in clinics are the most studied, contributing to more than 80% of entries (Fig. 1b). The drug targets of azoles and echinocandins, Cyp51 (also named Erg11 in yeast) and Fks, are unsurprisingly the most studied genes, being listed in nearly 40% and 25% of the entries, respectively. Although azoles and echinocandins are the most commonly used antifungals to treat invasive infection, this representation probably overestimates Cyp51 and Fks1 contribution to resistance. Indeed, many studies focus only on these genes as the potential cause of resistance and might falsely conclude that they found a resistance mutation or might miss a resistance mutation in another gene. The other genes are much less studied as about 90% of them are listed fewer than 25 times. Most of the biases in the entries can be explained by the fact that AMR in human fungal pathogens are overall better studied or more often reported than in plant pathogens. However, the diversity of species and antimicrobials reported for plant pathogens is larger than for human pathogens (Fig. 1b, c). Also, azoles are widely used in both clinics and agriculture, making them and their target Cyp51 the subject of many studies (Fig. 1b).

### Identifying AMR mutations and their impact on protein function

Antifungal resistance frequently arises through loss-of-function mutations (LOF). LOF mutations disrupt gene function, for instance, by introducing premature stop codons, InDels causing frameshift or missense mutations at critical sites in proteins. For some molecules, how LOF leads to resistance can be deduced from the antifungal mode of action. For instance, deletions of genes other than *CYP51* in the ergosterol biosynthesis pathway have been shown to confer resistance to azoles. These drugs bind to Cyp51, which blocks ergosterol biosynthesis and leads to the accumulation of a toxic dienol intermediate by the sterol desaturase Erg3^14^. The deletion of *ERG*3 prevents the synthesis of the toxic sterol^63^, causing resistance. Upstream of the pathway, the deletion of *HMG1*, an enzyme with sterol sensing functions, increases sterol production which causes resistance^63^. Another example is resistance to flucytosine (5-FC), a prodrug that must be converted by the cytosine deaminase to gain potency. A LOF in the gene encoding this enzyme (*FCY1* in *S. cerevisiae*) or other downstream genes that further process the molecule (e.g. *FUR1*) leads to resistance^64,65^.

We examined all instances of resistance associated with premature stop codons and InDels causing frameshifts to identify genes whose contribution to resistance may occur through LOF. LOF in *FCY1* and *FUR1* are well-established mechanisms of resistance in *S. cerevisiae* and unsurprisingly, they are also observed in pathogens such as *C. neoformans*^66^ and *N. glabratus*^67,68^. Experimental evolution of *S. cerevisiae* in nystatin led to the identification of LOF causing resistance in *ERG3* and *ERG6*^69^. Frameshifts and stop codons in *ERG3* and *ERG6* also appear to cause resistance to Amphotericin B in *C. auris*^70^, revealing that such LOF in these genes could be a conserved mechanism of resistance to polyenes. LOF in other genes of the ergosterol pathway is also associated with resistance, including frameshifts in *ERG2* in *C. albicans* causing resistance to azoles^71^. LOF of *ERG4* has also been associated with azole and echinocandin resistance in *N. glabratus*^72^ but with a confidence score of 8, which means that this requires further examination. Interestingly, all of these cases occur in species that are mainly haploid (or haploid strains of *S. cerevisiae*), which is anticipated as most LOF mutations are expected to be recessive.

Amino acid substitutions in Fcy1 or Fur1, typically at highly conserved sites, are often destabilizing for the protein and sufficiently disruptive to their function so that they also confer resistance^64,65^. It is therefore possible that conservation level and estimates of destabilization could help to better understand whether resistance emerges from LOF, without the need to observe premature stop codons or frameshift mutations in resistant isolates. To examine this and see if other mutations may be indicative that LOF causes AMR, we integrated into the data two predictions of the impact of mutations based on conservation analysis^24,73^ and one based on the prediction of the destabilization by the mutation using protein structure^25,26^ (Fig. 2). For genes that lead to resistance upon LOF, such as *FCY1* and *FUR1*, we indeed clearly see that resistance mutations are predicted to have a larger impact on protein function and stability in comparison to other mutations in the same genes. Other extreme cases are Sur1 and Csg2, proteins involved in sphingolipid synthesis. Amino acid substitutions in these proteins that cause resistance are particularly destabilizing and occur at conserved sites, suggesting LOF. This observation is validated by recent experimental evolution results showing that stop codons in these two enzymes in *S. cerevisiae* cause resistance to clotrimazole^74^. For *ERG3* and *HMG1*, mutations are predicted to have a greater impact on the protein function but to have a low or no impact on protein stability (*ERG6* did not have enough entries to perform this analysis). Nonetheless, this suggests that resistance mutations in *ERG3* and *HMG1* likely arose through LOF which can be predicted through their potential effects on protein function.

**Fig. 2.**
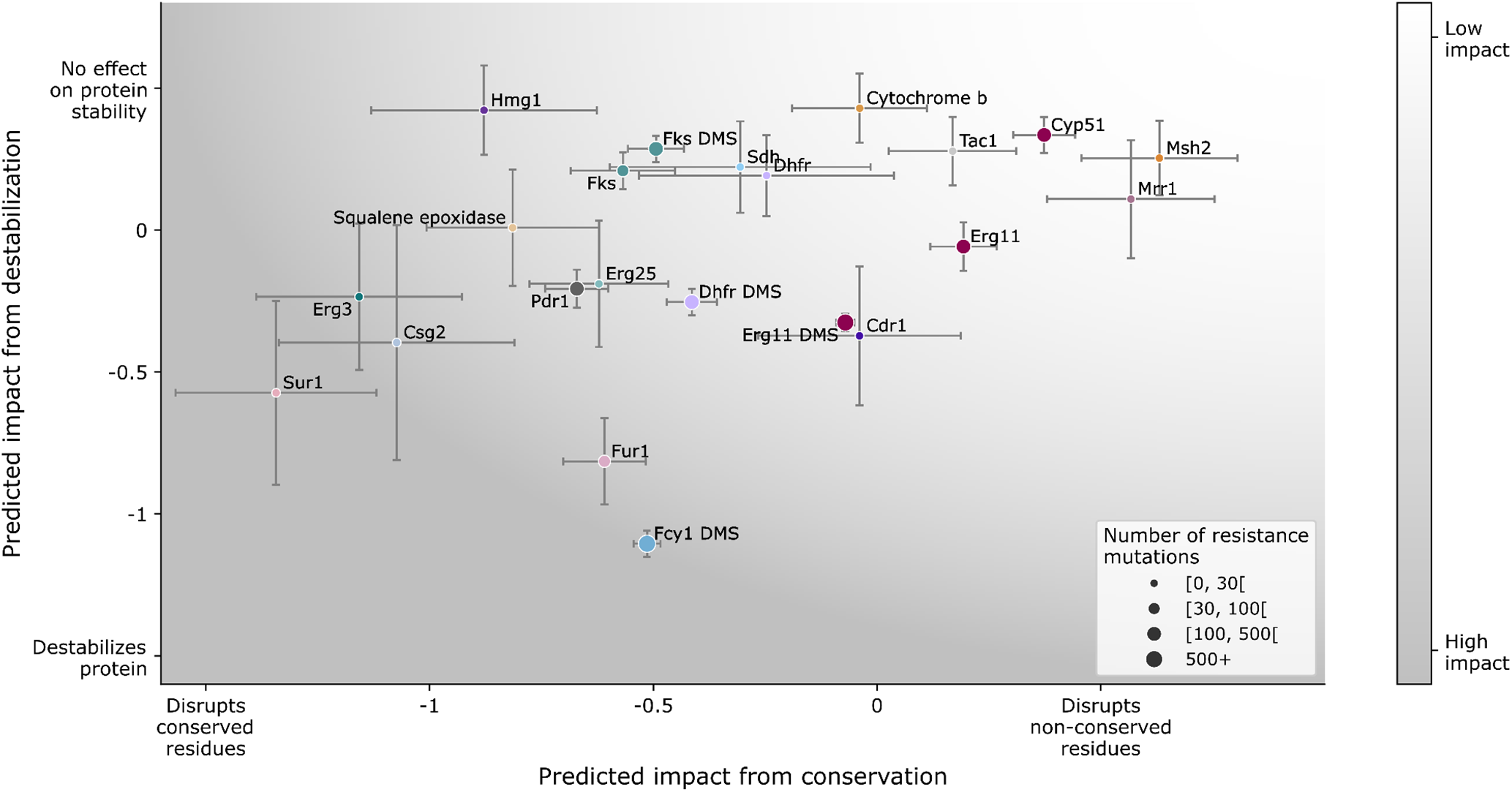
Prediction of mutations’ effect on protein function and stability can aid differentiation between loss and gain of function resistance mutations. Relationship between the predicted impact of resistance mutations inferred from long-term evolutionary constraints (GEMME^24^) and protein stability (FoldX^25,26^). Each dot represents the mean deviation of the predicted impact for resistance mutations in a given protein relative to all possible amino acid changes. The y-axis presents the z-score of the normalized predicted change in the Gibbs free energy (ΔΔG) upon mutation predicted by FoldX. Negative values predict destabilization of the protein. The x-axis represents the z-score of the predicted impact of the mutation computed by GEMME, which uses patterns of protein conservation to predict the tolerance of mutations. Negative values predict a higher impact on the protein function based on evolutionary conservation. For each gene, when mutations were available from DMS experiments (confidence score of 3), the DMS data is shown separately. The size of the dots shows the sample size and the colors show the different proteins. Error bars represent one standard error. The distribution of the data for each gene is presented in Extended data Fig. 1.

As most resistance genes for which we have enough data to perform such analysis do not show patterns consistent with LOF mutations, this suggests that a large fraction of resistance genes does not evolve by simple inactivation. In fact, antifungal resistance can also arise through gain of function (GOF) mutations. The mutational mechanisms by which a gene gains a function by mutations can be diverse (e.g. premature stop codons, InDels or missense mutations). One way GOF mutations can lead to resistance is through the production of an hyperactive protein. For instance, mutations in the transcription factor Upc2 increase the expression of the azole drug target, which compensates for the inhibition by azoles^75^. Alternatively, GOF mutations in efflux pumps such as Cdr1/Pdr5 and Mdr1^76–78^ or in their transcription factors such as Tac1, Pdr1 and Mrr1^78^ increase drug efflux outside the cells, and are often responsible for multi-drug resistance. Another more ambiguous type of GOF are resistance mutations in some drug target proteins, such as Cyp51 for azoles or Fks1 for echinocandins, that can prevent drug binding while maintaining the regular activity of the enzymes^79^. Although the loss of drug binding could be the consequence of a LOF phenotype, it cannot lead to drug resistance if the natural substrate cannot be processed. As expected for GOF resistance mechanisms, the impact of resistance mutations on protein function and stability are predicted to be lower than for LOF resistance mutations (Fig. 2). Consequently, GOF mechanisms are more difficult to identify directly from the predicted impact of mutations on the proteins.

### Mapping AMR mutations on protein structures

Mutations leading to LOF through destabilization need to be located in regions of proteins important for stability. Alternatively, they could occur at the active sites of the proteins. To help rationalize how mutations may confer resistance, we colored residues with listed resistance mutations on the protein structures to detect potential clustering. Fur1 and Fcy1 are tetrameric and dimeric proteins, respectively. In both cases, many of the mutations occur in structured and core residues, including at the interfaces, potentially disrupting complex assembly (Fig 3a, b). Erg3 and Hmg1 structures were predicted with AlphaFold 3^27^. For Erg3 (Fig. 3c, pTM = 0.91), reported mutations are in transmembrane helices. For Hmg1, Alphafold 3 predictions have a lower confidence level (Fig. 3d, ipTM = 0.64 and pTM = 0.47), but prior knowledge on the protein allows us to deduce that mutations are located in the conserved sterol sensing domain of the protein^80^. This could explain the results above (Fig. 2) where resistance mutations occur at conserved sites but are not predicted to destabilize the protein structure. Although the structures presented are not models of how mutations impact the structure of the proteins, the mapping can help interpret the impact of mutations.

**Fig. 3.**
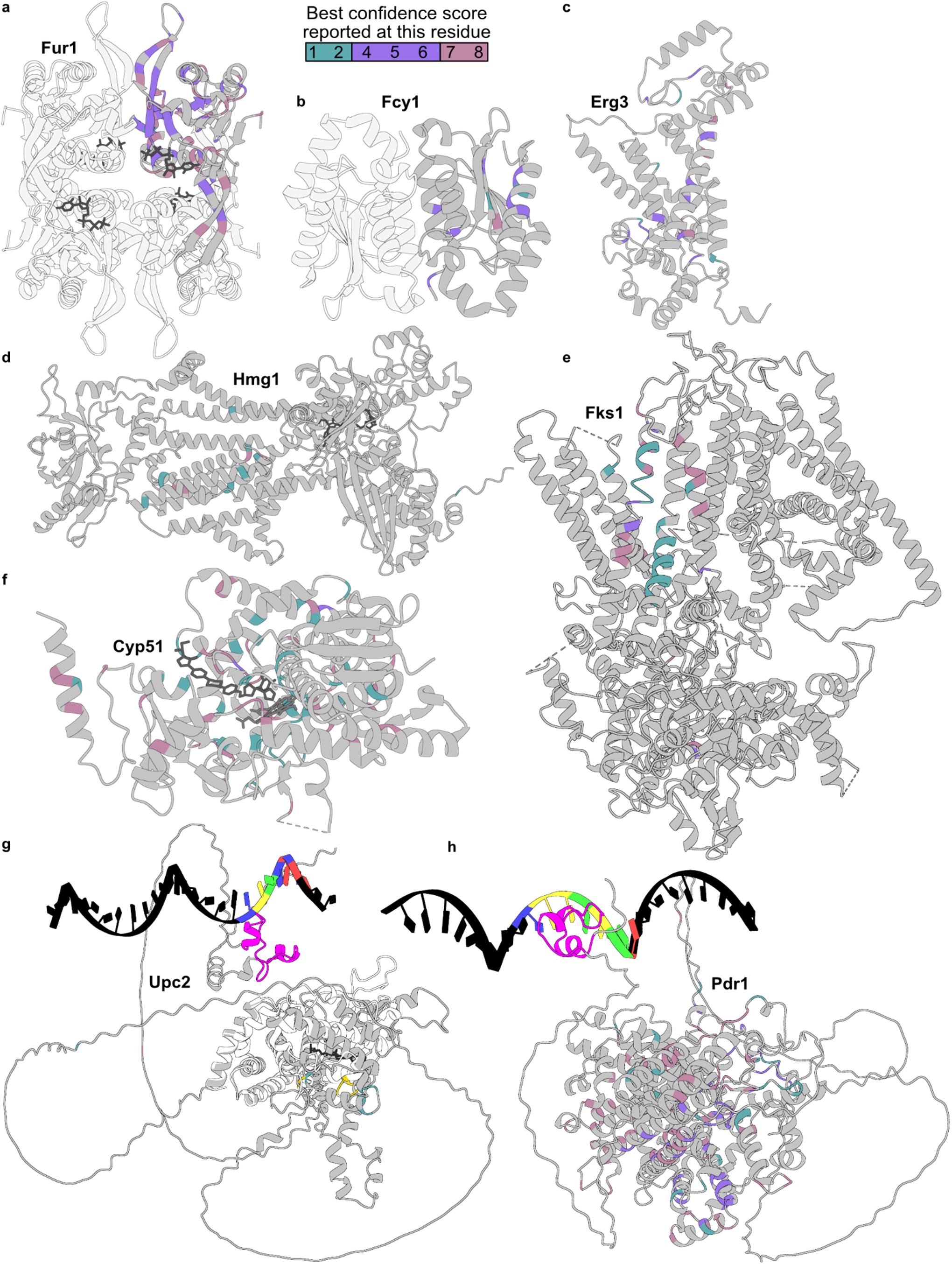
Mapping of resistance mutations on protein tertiary structures. **a,b,c,d,e,f,g,h** For each protein structure, residues are colored by the best positive confidence score for all reported mutations at this residue. Multiple sequence alignments were used to map resistance mutations of orthologs. Confidence scores of 3 (DMS assays) were excluded. *C. albicans* Fur1 in complex with UTP (PDB ID 7RH8) (**a**), *S. cerevisiae* Fcy1 (PDB ID 1P6O) (**b**), *S. cerevisiae* Erg3 (AlphaFold 3 prediction with Uniprot P32353 protein sequence, pTM = 0.91) (**c**), *A. fumigatus* Hmg1 (AlphaFold 3 prediction with Uniprot Q4WHZ1 protein sequence and NADP, ipTM = 0.64 and pTM = 0.47) (**d**), *S. cerevisiae* Fks1 (PDB ID 7XE4) (**e**), *C. albicans* Erg11 bound to itraconazole (PDB ID 5V5Z) (**f**), *C. albicans* Upc2 (AlphaFold 3 prediction with Uniprot Q59QC7 protein sequence and AATATCGTACCCGATTATG**TCGTATA**TT DNA sequence^28^, ipTM = 0.37 and pTM = 0.45, grey) superposed with *N. glabrata* Upc2 ligand-binding domain in complex with ergosterol (PDB ID 7VPR, white with orthologous resistance mutations in dark yellow) (**g**), *S. cerevisiae* Pdr1 (AlphaFold 3 prediction with Uniprot P12383 protein sequence and GAGAAATGTC**TCCGCGGA**ACTCTTCTAC DNA sequence^30^, ipTM = 0.4 and pTM = 0.69) (**h**). **g,h** In bright pink are the Zn2-C6 fungal-type DNA binding domains. On the DNA strands, colors represent the nucleotides (T:Blue, C:Yellow, G:Green, A:Red) of the core Upc2 DNA binding site **TCGTATA** of *C. albicans ERG11* promoter^28,29^ (**g**) and of the pleiotropic drug response element (PDRE) **TCCGCGGA** present in *S. cerevisiae PDR5* promoter^30^ (**h**).

Mutations involved in preventing drug binding are expected to cluster in regions of proteins where the drug and the protein interact. We mapped resistance mutations on the protein structures for proteins that are the most represented in the literature. For Fks, echinocandin resistance mutations are clustered in three different hotspots (Fig. 3e) as it has been previously observed^81,82^. However, there are also reported resistance mutations outside of the described hotspots, suggesting they might be larger than previously proposed (although we note some of these mutations have low confidence scores). The proximity of the hotspots on the enzyme structure suggests a putative binding site of echinocandin drugs^82,83^. For Cyp51, azole resistance mutations are more dispersed. However, some regions of the protein appear to be enriched with such mutations (Fig. 3f). A DMS of *C. albicans* Cyp51 (Erg11) has been conducted, and the authors similarly concluded that resistance mutations are dispersed and not limited to distinct clusters^84^.

One potential way to identify GOF mutations in transcription factors would be to examine whether mutations cluster in space in specific domains or regions associated with particular functions. No complete experimentally determined structures were available for any lead transcription factors. Therefore, we predicted two of them using Alphafold 3^27^ in order to map resistance mutations. The prediction scores are low (Upc2 : ipTM = 0.37 and pTM = 0.45, Pdr1 : ipTM = 0.4 and pTM = 0.69), meaning we cannot be confident of the predicted structure. For Upc2, the structure of the ligand-binding domain in complex with ergosterol has been determined experimentally. Therefore, we superposed the crystalized ligand-binding domain with the complete Alphafold 3 predicted structure (Fig. 3g). We found that the few reported resistance mutations cluster in Upc2 but not in the DNA binding region. Mutations in Upc2 have already been found to increase the protein activity through the interruption of ergosterol binding in the sterol-binding pocket^85^. For Pdr1, although the predicted structure is of low confidence, we see that the resistance mutations are located across the structure, but not at the DNA binding site (Fig. 3h). There is evidence that GOF mutations in Pdr1 do not result in greater DNA binding activity, but instead block inputs that would normally limit Pdr1 transcriptional activation^86^. Interestingly, Pdr1 mutations are in the middle range in terms of predicted effects based on conservation and destabilization (Fig. 2), suggesting that mutations could modify Pdr1 activity through minor destabilization. For example, Pdr1 with GOF mutations in *N. glabratus* have been shown to be less stable than the wild-type^87^. These observations from Upc2 and Pdr1 suggest that resistance mutations in transcription factors might often not directly alter DNA binding but rather other facets of their activity.

### Extreme convergence of AMR mutations among species

The use of the same antifungals to treat infections by different species allows us to examine evolutionary convergence in the mechanisms of resistance. Convergence is often used as evidence that a trait has been under selection, which in the case of AMR helps interpret and prioritize mutations for validation and for modifying drugs to overcome frequently encountered resistance mutations^88^. Experimental evolution studies have shown that the diversity of mutated genes to adapt to antifungals was narrow within a species and often overlapped with mechanisms described in other fungi^69,89,90^. Convergence in the molecular bases of resistance could occur at several levels: in the orthologous genes at the same sites (mutational level); in orthologous genes but at different sites (gene level); or at the functional level, involving genes with similar functions but that are not homologous. We first examined the level of convergence in the resistance mechanisms among fungi at the gene and at the functional levels by comparing resistance genes between species (Fig. 4a, b, Extended data Fig. 2). Outside of the specific drug targets such as Erg11 and Fks, which are expected to be convergent, we found that, generally, fungi evolve resistance through mutations in homologous genes or in genes with similar functions. One example of this is efflux pumps of the ABC (e.g. Cdr1 of *C. albicans* and Mdr2 of *A. fumigatus*) and MFS (e.g. Mfs1 of *A. flavus*) transporter families^78^ or transcription factors that regulate efflux pumps expression (e.g. Tac1 of *C. albicans*, Pdr1 of *N. glabratus* and Mrr1 of *C. albicans*). Resistance also results from variations in the number of copies of genes ^91^. However, these copy number variations often have the same effects as point mutations. For example, azole resistance can arise from *ERG11* overexpression caused by a gene copy number increase ^92^. The absence of convergence, or species-specificity in AMR mechanisms, is more difficult to assess due to incomplete data resulting from varying research efforts across species.

**Fig. 4.**
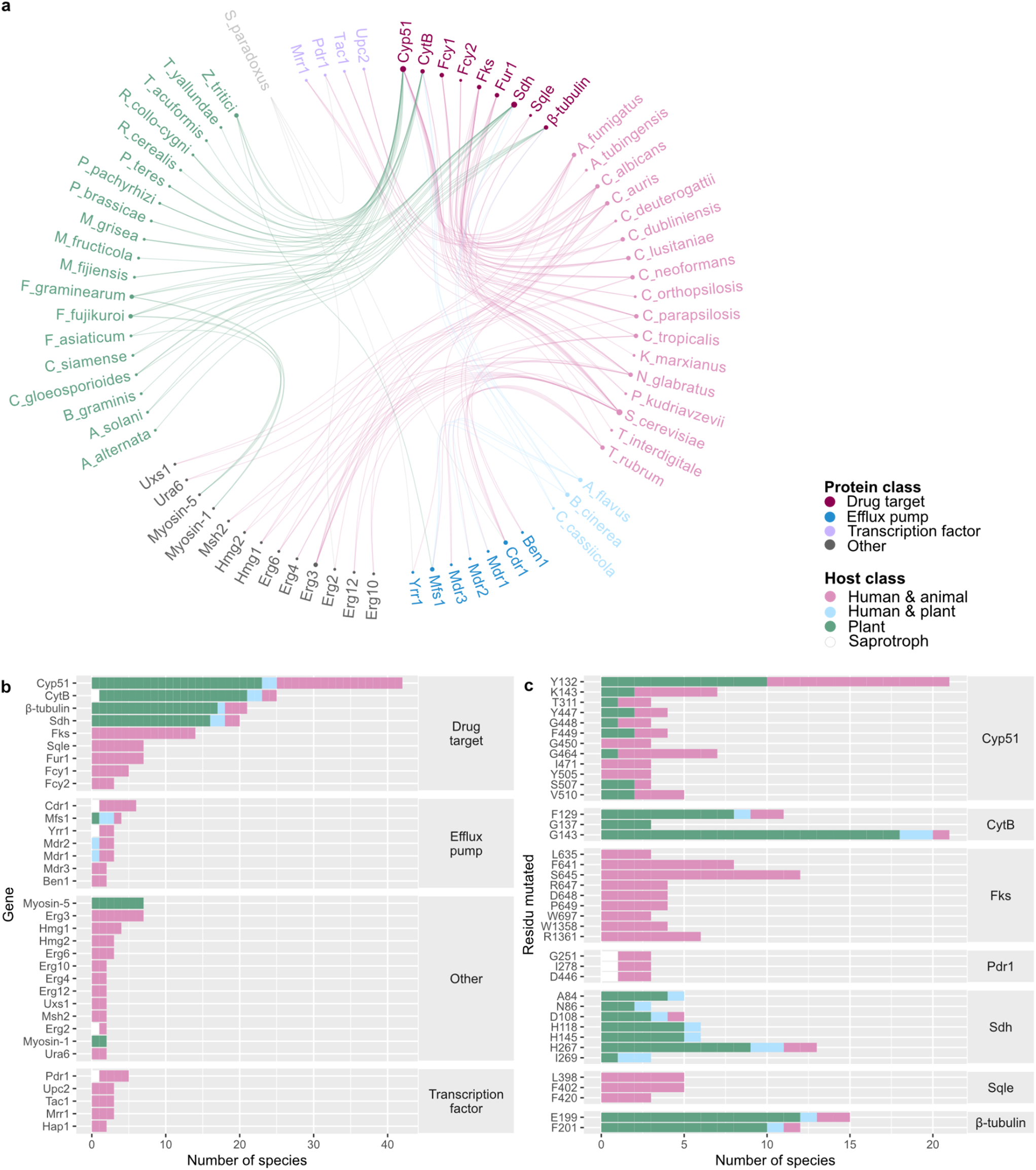
Massive convergence in the mechanisms of resistance among fungi at the gene and mutation level. **a,** Edgebundle plot^99^ of mutated proteins associated with resistance in more than one fungal pathogen. Only proteins and species with more than five entries were included in this analysis. The interactive html file is available in Supplementary data 2. Genes classes were colored based on their function. **b,** Number of species with the same mutated protein associated with resistance. Only instances of proteins associated with more than two species are shown for clarity. **c,** Orthologous mutations conferring resistance in several species show a high level of convergence at the mutation level. Cyp51, Sqle (Squalene epoxidase) and Fks use *C. albicans* numbering in the MSA. SdhB, SdhC, SdhD, Beta-tubulin and Cytb (Cytochrome b) use *Z.tritici* numbering in the MSA. Pdr1 uses *S.cerevisiae* numbering. Only instances of mutations associated with more than two species are shown for clarity. Entries with multiple proteins were excluded.

Beyond gene-level convergence, we can also examine convergence at the level of individual mutations. Some proteins, such as Cyp51 and Fks, are well conserved across fungi, and it is therefore possible that the same mutations would confer resistance across species. Using multiple sequence alignments (MSA), we compared the identity of resistance mutations across orthologs. We found several cases where the same resistance mutations occur in two or more species (Fig. 4c). The most impressive examples are amino acid substitutions in Cyp51. For instance, Y132H and F in Cyp51 (*C. albicans* aa numbering) have been found to confer azole resistance in more than 20 species, including the distantly related species *C. albicans, A. fumigatus and C. neoformans*. Amino acid substitutions at position Y132 have been found to disrupt azole binding^93^. Other striking examples are mutations at position S645 of Fks (*C. albicans* aa numbering) and G143 of the cytochrome b (*A. fumigatus* aa numbering), which are associated with resistance in 10 species or more. In Fks, S645 is located in the echinocandin resistance hotspot 1 and is most likely involved in drug binding^82,83^. Mutations at the residue S645 have been reported to decrease the maximum velocity of Fks and have been linked to increased echinocandin clinical failure^94–97^. For cytochrome b, it is hypothesized that the G143A mutation alters the binding of Qo inhibitors, which are a class of fungicides that target the quinol oxidation (Qo) site of the cytochrome bc1 complex, through steric interactions^98^. Mutations at residues where resistance mutations with a high level of convergence are observed can confidently be interpreted as resistance mutations across species. Such convergence could be explained by mutation bias, but given the breadth of species concerned, it is likely that convergence is rather due to the fact that these mutations confer high resistance levels with little to no evolutionary tradeoff in comparison to other possible resistance mutations^88^. We provide the MSA of the most frequently reported genes across multiple species as a resource (Supplementary data 1).

### Cross-resistance is common between and within classes of antifungals

One major question regarding AMR mutations is whether they confer cross-resistance to more than one drug, either of the same class or different classes. FungAMR allows quantifying the extent of cross-resistance. We found that if a mutation in a gene provides resistance to one antifungal, it will likely confer cross-resistance to others within a class compared to between classes (Fig. 5a, b, Extended data Fig. 2). This was expected given the similarities of molecules within a class (Extended data Fig. 3) and their shared mode of action^100^. For instance, we found extensive cross-resistance between clinical and agricultural azole antifungals. However, cross-resistance is not a systematic consequence of mutations given the differences in the chemical structures of the drugs (Extended data Fig. 3). Since not all mutations present in our dataset were systematically assayed for multiple drugs, it is difficult to identify clear cases of resistance mutations that are specific to only one drug. Our extensive data nevertheless provides a global view of which mutations in which genes confer cross-resistance within and between drug classes.

**Fig. 5.**
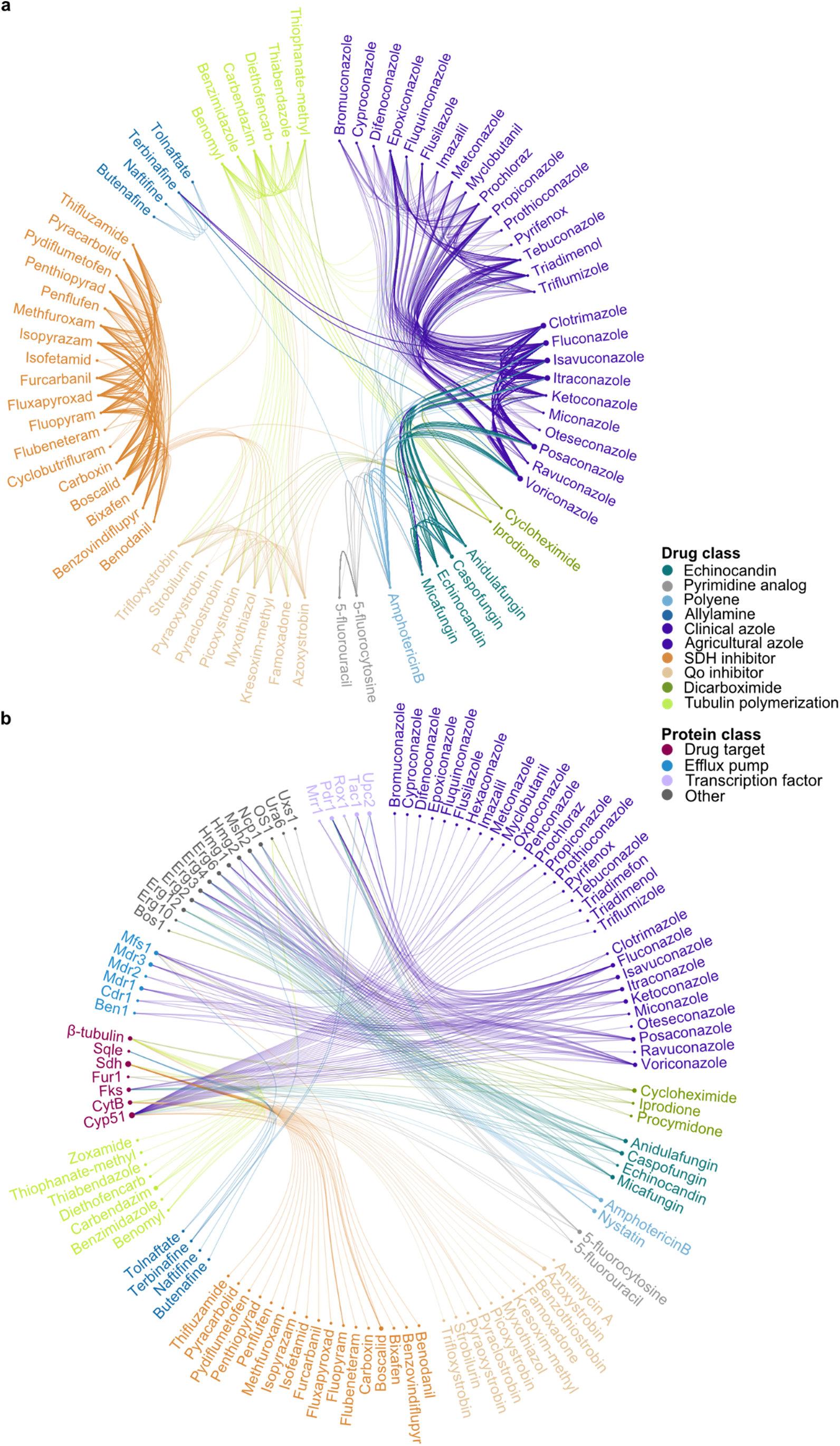
Rampant cross-resistance between and within classes of antifungals. **a,** Edgebundle plot^99^ of cross-resistance between antifungals. Drugs are linked when a unique mutation in a single protein can confer resistance to multiple drugs. Only proteins and drugs with more than five entries were included. The interactive html file is available in Supplementary data 3. **b,** Edgebundle plot^99^ of mutated proteins associated with at least one mutation conferring resistance to more than one clinical or agricultural antifungal. Only proteins and drugs with more than five entries were included. Entries with multiple proteins mutated in the same strain were excluded. The interactive html file is available in Supplementary data 4.

Although within-class cross-resistance is more frequent, between-class cross-resistance also occurs^101,102^. Mutations found in clinical isolates in proteins implicated in ergosterol biosynthesis such as Cyp51, Upc2, Hmg1, Erg3 and Erg6 can confer resistance to both azoles and polyenes, even if these antifungals target different elements of the ergosterol biosynthesis pathway (Fig. 5b). Additionally, some mutations in Fks or Erg proteins seem to be involved in both azole and echinocandin resistance (Fig. 5b). This connection is more surprising since echinocandins do not target cell membrane proteins but target a beta-glucan synthase, and beta-glucan is a component of the fungal cell wall. Studies have shown that changes in the cell wall or membrane composition can provide an advantage for cells under drug-induced stress^103–105^, possibly making resistance to the different clinical antifungal classes more interconnected than anticipated from the drugs’ mode of action. Cross-resistance caused by interconnection of transcriptional pathways has been reported before^90,106106^, but our observations suggest that cross-resistance between classes could also happen from mutations in the drug targets. On the side of agricultural antifungals, cross-resistance between the different classes is also present, but seems to be less frequent than among human pathogens. Such a result could also be due to lack of data, as resistance has been less frequently tested systematically among classes of plant antifungals. Although incomplete, these analyses are sufficient to further strengthen important concerns. First, cross-resistance between medical and agricultural azoles is frequent. Second, azoles, polyenes and echinocandins are connected through a core set of genes, meaning that cross-resistance to clinical drugs can evolve through single mutations. The full extent of antifungal cross-resistance remains to be assessed but this analysis reveals that three out of the four classes of drugs used to treat fungal infections are not entirely orthogonal. This interdependence stresses the need for finding new molecular targets and drugs for treatment and most importantly, increase stewardship between clinical and agricultural drugs when new drugs are developed and new targets identified^102^.

### ChroQueTas efficiently screen AMR in fungal genomes

To facilitate the analysis of AMR in fungal genomes from the FungAMR data, we developed ChroQueTas, a user-friendly bioinformatic tool that only necessitates a fungal genome in FASTA format. As a practical example of how ChroQueTas can be used to identify AMR mutations using FungAMR, we downloaded the raw sequencing data (Illumina) from 46 *C. albicans* and 144 *Zymoseptoria tritici* isolates^107–109^ (Supplementary table 5). These two species were selected to represent human and agricultural pathogens, respectively. The *C. albicans* dataset included isolates from two countries (China and the United States of America). The *Z. tritici* dataset included isolates recovered from wheat from four countries (Australia, Israel, Switzerland and the United States of America). After quality control of the sequencing data and further assembly, the genomes were submitted to ChroQueTas by choosing their species-associated scheme, which performed AMR screening over all the target proteins described in FungAMR for these two species, and we report here the results for Cyp51 (Erg11) due to their key association with AMR.

The screening of the *C. albicans* and *Z. tritici* genomes performed with ChroQueTas was based on FungAMR data, which contains the largest number of AMR mutations reported to date, and thus, we were able to identify all the AMR mutations reported by the authors. As a summary, we found that 83.3% of the selected *C. albicans* genomes harbored at least one mutation in Erg11 known to cause AMR (Fig. 6a). The most commonly identified mutated positions were A114 (24% of genomes, n=11) and Y132 (24% of genomes). Nine of the *C. albicans* that harbored the mutation A114S also harbored Y257H, a combination that has been previously reported^110^ to cause AMR against different azoles, including fluconazole and itraconazole. The other two genomes (SRR13587870 and SRR13587591) harbored mutation A114V (previously reported to not cause AMR alone^84^) together with F145L, a combination that have shown to confer resistance against fluconazole^111^. Regarding the position Y132, the nine Chinese isolates harbored the mutation Y132H, whereas the two U.S. isolates harbored the mutation Y132F, also associated with azole resistance. For the analysis of the *Z. tritici* genomes, 84.7% of the isolates showed at least one mutation in Cyp51 known to confer resistance (Fig. 6b). Additionally, 30.5% of the genomes (n=44) harbored Y459Del and/or G460Del deletions, associated with AMR against different azoles^112,113^. Additionally, the combination of ChroQueTas and FungAMR allowed us to identify eight additional AMR mutations in *C. albicans* and one in *Z. tritici* that were not reported by the authors in their corresponding studies (highlighted in green in Fig. 6). Additionally, we identified Y461D in SRR3740350 (*Z. tritici* isolated in Switzerland) which occurs at a position previously associated with AMR due to another mutation, suggesting this mutation may also contribute to resistance. The results summarized here show the strength of ChroQueTas and FungAMR to efficiently screen AMR in fungal genomes and the potential to identify novel mutations involved in AMR.

**Fig. 6.**
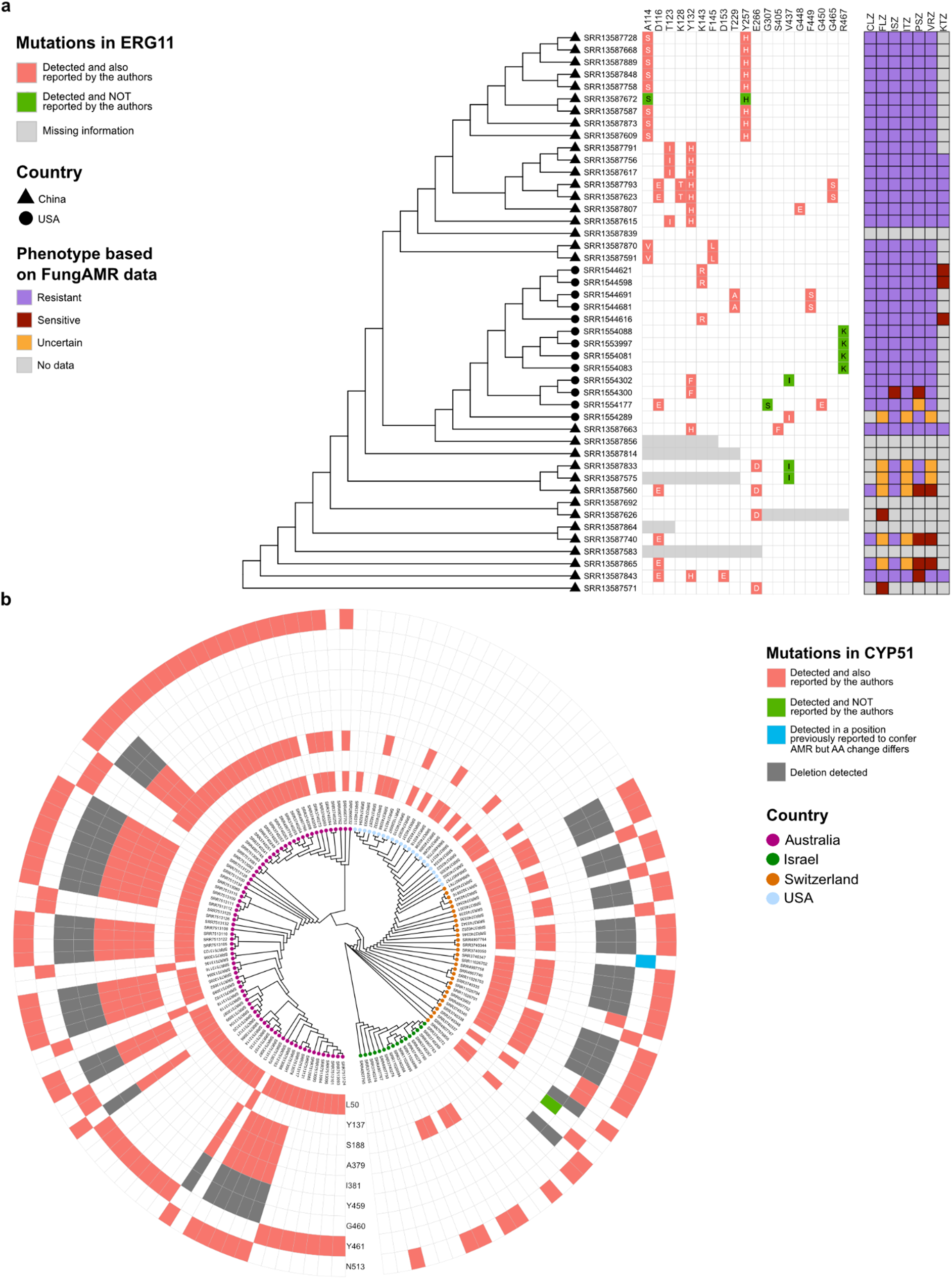
Practical example of AMR screening by using ChroQueTas and FungAMR. **a, b,** Phylogenetic comparison of *C. albicans* (**a**) and *Z. tritici* (**b**) datasets based on average nucleotide identity (ANI). The trees’ tips are coloured or shaped according to the country in which they were isolated. Each heatmap represents the position within the query protein sequences (Erg11 for *C. albicans* and Cyp51 for *Z. tritici*) and are coloured when a mutation was detected with ChroQueTas. AA stands for amino acids. “Missing information” in (**a**) stands for proteins that could only be resolved partially due to genome fragmentation and “Uncertain” refers to mutations that are associated with a low positive and a high negative evidence scores making the strain phenotype uncertain CLZ, clotrimazole; FLZ, fluconazole; ISZ, isavuconazole; ITZ, itraconazole; PSZ, posaconazole; VRZ, voriconazole; and KTZ, ketoconazole.

## Discussion

Despite the threat that fungal pathogens and AMR represents, we were still missing a high quality centralized antifungal mutations repository. Here, we present FungAMR, a compendium of 35,792 carefully curated entries of clinical and agricultural importance. One major finding is that many resistance mutations reported in the literature have little experimental support beyond identifying a mutation in a known resistance gene in a resistant isolate, which can be coincidental (confidence scores of 7 and 8, Fig. 1a). Such low quality reports create an overconfidence in our knowledge of the identity of resistance mutations. Our observations call for more experiments and for the development of computational tools focusing on the direct association of mutations to resistance. The reconstruction of individual mutations in pathogens will always remain more challenging than in model organisms but recent progress with genome editing directly in the pathogens makes such experiments possible on an appreciable scale (confidence score of 1, Fig. 1a). Heterologous expression of resistance genes in a model organism such as *S. cerevisiae* has been proven to be a powerful solution in a large number of studies (confidence score of 2, Fig. 1a). Since human and *E. coli* genes^114,115^ can complement many of *S. cerevisiae*’s orthologs, it is likely that most genes present in FungAMR can be expressed and be studied in model species. Furthermore, high throughput experiments such as deep mutational scanning are powerful ways to characterize a large number of genotype-resistance phenotype associations^64,79,84,116,117^(confidence score of 3, Fig. 1a)^64,79,84,116,117^. As we show, the fact that AMR mutations share specific features and locations in proteins also opens the possibility of using computational approaches to further increase confidence into the likely causal role of mutations that could otherwise not be validated experimentally.

FungAMR also highlights how research is extremely skewed toward a few fungal species, genes and drugs (Fig. 1). The scientific community would benefit from a diversification of studies. For instance, by focusing analysis on only a few genes that are *a priori* known to be associated with resistance, we are limiting our comprehension of the full suite of different resistance mechanisms. Our ability to screen for resistant isolates through whole genome sequencing should greatly facilitate such an approach, yet sometimes even such datasets are only mined for variants in a subset of genes. Also, for most genes, our knowledge of resistance mutations across different species is too limited to assess how convergent they are. The same applies for the impact of mutations on AMR within and between classes in order to assess the level of cross-resistance. Although limited, the information we currently have is crucial in our fight against fungal pathogens and suggest that there is a high level of convergence in the mechanisms of resistance among fungi (Fig. 4) and that cross-resistance seems to be common within and between classes of antifungals (Fig. 5). FungAMR has the potential to help predict resistance mutations and develop new drugs, but more studies incorporating different species and different antifungals are needed to determine how universal our observations are. In the meantime, there is still an urgent need to identify new molecular targets and drugs for treatment, and the dual use of some antifungal classes in the clinic and in agriculture should be considered a critical threat to our ability to treat human fungal infections. Fisher et al.^118^ propose a roadmap with actionable guidance for researchers to better understand and manage fungal antimicrobial resistance across one health.

In addition to providing an AMR mutation catalog, we show that variant effect predictors based on protein stability and sequence conservation can be used to differentiate between LOF and GOF resistance mechanisms. Along with structural mapping, these tools can be used to interpret the impact of mutations in proteins involved in antifungal resistance. However, we are missing structural models for most proteins listed in FungAMR. Even though advances are made in protein structure predictions, algorithms such as AlphaFold are dependent on training data availability. Even if predictions are extremely valuable to interpret the impact of mutations in a protein, accurate knowledge of the 3D structure is crucial for protein structure-based drug design^119^. The scientific community would greatly benefit from experimentally determined structures of proteins associated with resistance.

Futhermore, we developed ChroQueTas, a user-friendly bioinformatic tool based on the FungAMR resource, to facilitate the screening of AMR mutations in fungal genomes. We show that ChroQueTas and FungAMR together allow efficient screening of AMR in published fungal genomes by identifying all resistance mutations reported by the authors in addition to new resistance mutations that were not reported initially.

Finally, it is important to note that FungAMR is intended to be a general resource to better understand resistance mechanisms rather than a guide to predict treatment outcomes. FungAMR data are based entirely on diverse in vitro assays reported by hundreds of researchers in the literature, which may not directly reflect clinical or field observations. In addition, treatment success or failure is influenced by several factors beyond the scope of this work, such as host immune responses, drug pharmacodynamics, biofilm formation and pathogen strain variability^10,120,121^. Therefore, although in vitro susceptibility testing is useful for identifying isolates less likely to respond to a drug, it may not correlate with the outcome of an antifungal treatment ^122^. Future studies will be needed to link in vitro susceptibility testing and the complex parameters in infected hosts to potential treatment outcomes^10^. Nonetheless, comprehensive datasets such as FungAMR are an important step towards this objective^120^.

## Acknowledgements

We thank Andrew McArthur and his team from McMaster University for setting up FungAMR as a web-searchable interface and for hosting it on the CARD database website.

## Data availability

The FungAMR resource is available in Supplementary table 1 and in a web-searchable interface on the CARD database website (https://card.mcmaster.ca/). The most up-to-date version of FungAMR can always be found on GitHub at https://github.com/Landrylab/FungAMR. The multiple sequence alignment files for the genes present in FungAMR are available in Supplementary data 1. The dataset used for the Supplementary note 1 is available at 10.5281/zenodo.12583470. The Protein Data Bank is available at https://www.rcsb.org/. UniProt is available at <u>UniProt</u>. The AlphaFold database is available at <u>AlphaFold Protein Structure</u> <u>Database</u>.

## Code availability

All scripts for figures and for FungAMR content analysis are available on GitHub at https://github.com/Landrylab/FungAMR. ChroQueTas is available on GitHub at https://github.com/nmquijada/ChroQueTas and Conda at https://anaconda.org/nmquijada/chroquetas.

## Extended data and figures

**Extended data Fig. 1.**
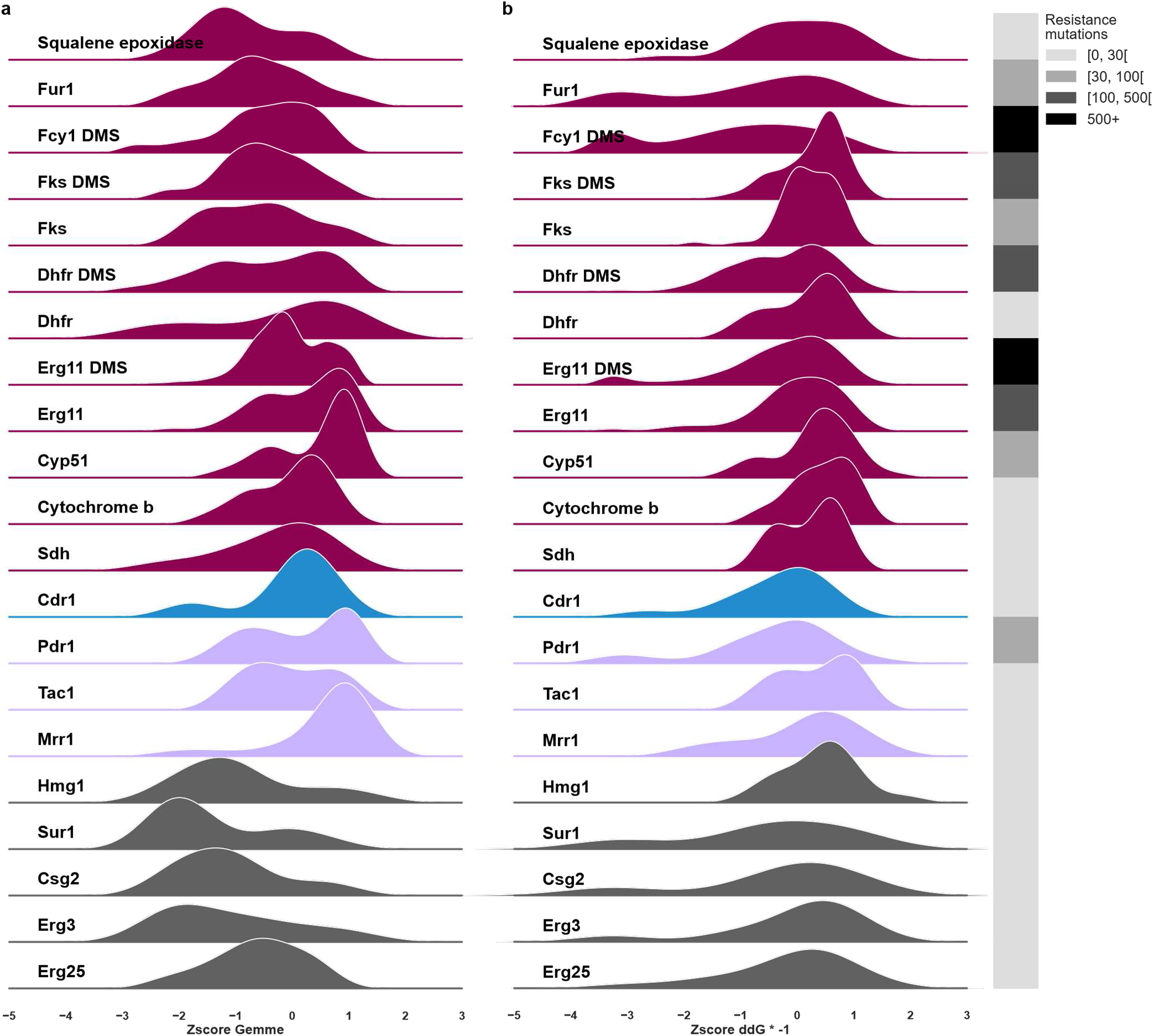
Distribution of predicted effect of AMR mutations on protein function and stability. **a,** Distribution of the standardized GEMME score for all unique resistance mutations across proteins. Negative values predict a higher impact on the protein function based on evolutionary conservation. **b,** Distribution of the standardized negated predicted change in the Gibbs free energy (ΔΔG) upon mutation predicted by FoldX for all unique resistance mutations across proteins. Negative values predict destabilization of the protein. For each protein, when mutations were available from DMS experiments (confidence score of 3), the DMS data is shown separately. The grayscale shows the number of unique resistance mutations per protein.

**Extended data Fig. 2.**
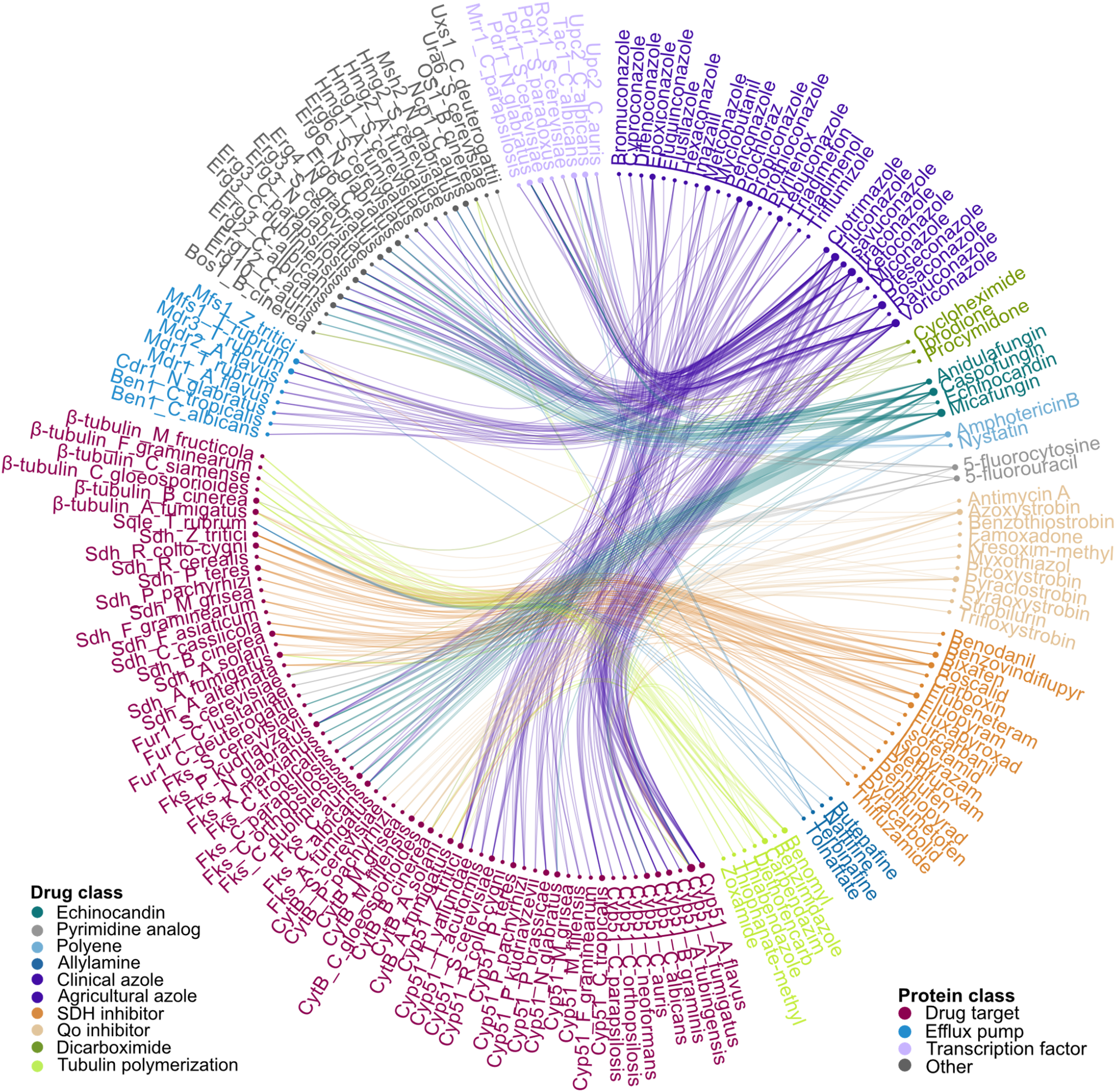
Convergence of resistance mechanisms and cross-resistance between antifungals. Edgebundle plot^99^ linking species specific proteins to antifungal resistance. Orthologous proteins are associated with drug resistance across multiple fungal pathogens, demonstrating evolutionary convergence. Furthermore, mutations in the same protein are linked to resistance against multiple drugs and drug families, indicating cross-resistance. Only proteins and drugs with more than five entries were included. The interactive html file is available in Supplementary data 5.

**Extended data Fig. 3.**
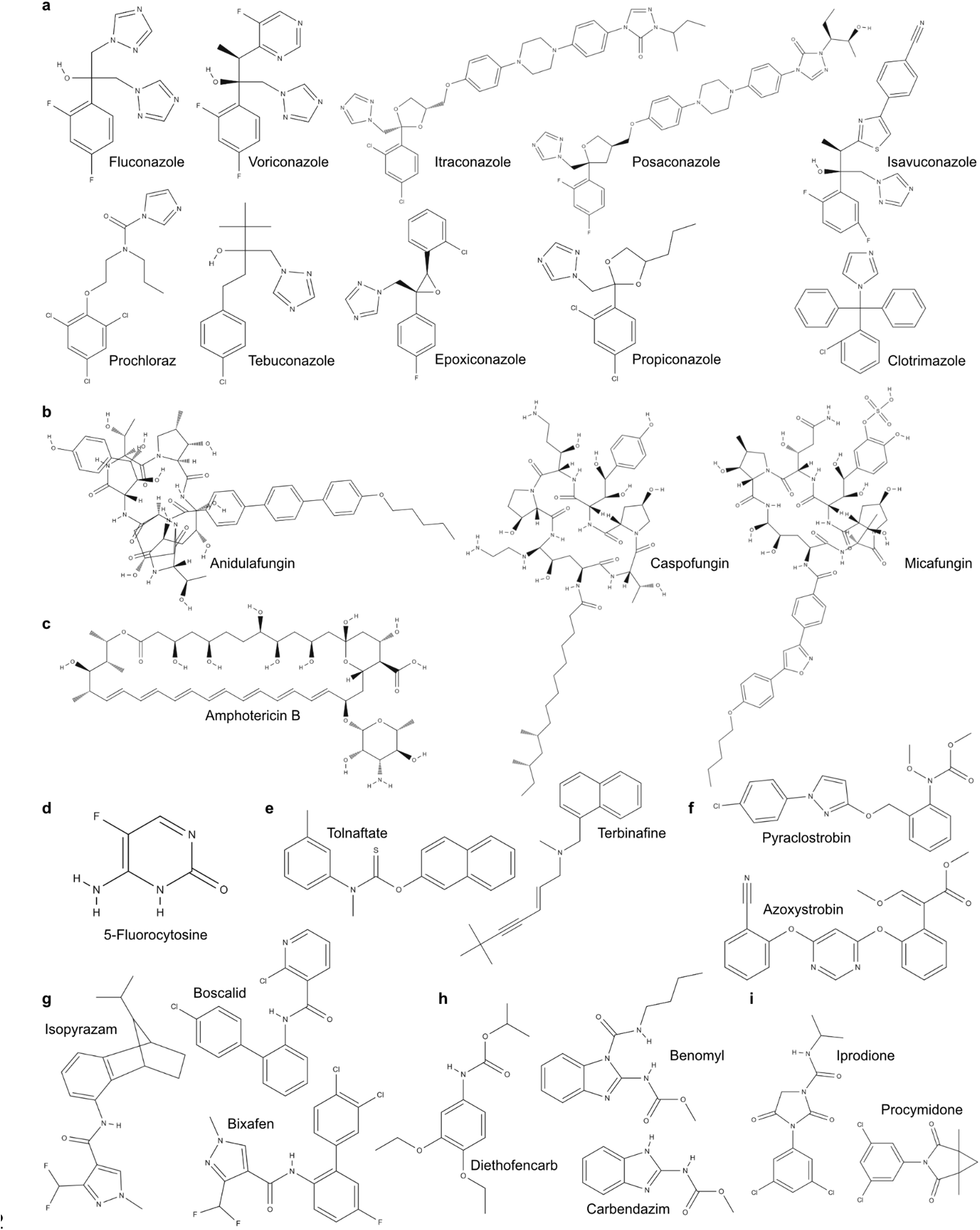
Chemical structures of antifungals from different classes. **a,** Fluconazole, voriconazole, itraconazole, posaconazole and isavuconazole are triazoles used in clinics. Tebuconazole, epoxiconazole and propiconazole are triazoles used in agriculture. Clotrimazole and prochloraz are imidazole used in clinics and in agriculture respectively. **b,** Anidulafungin, caspofungin and micafungin are echinocandins used in clinics. **c,** Amphotericin B is a polyene used in clinics. **d,** 5-Fluorocytosine (5-FC) is a pyrimidine analog used in clinics. **e,** Tolfanate and terbinafine are allylamines used to treat topical infections. **f,** Pyraclostrobin and azoxystrobin are quinone outside inhibitors (QoI) used in agriculture. **g,** Isopyrazam, boscalid and bixafen are succinate dehydrogenase inhibitors (SDHi) used in agriculture. **h,** Diethofencarb, carbendazim and benomyl are inhibitors of tubulin polymerization used in agriculture. **i,** Iprodione and procymidone are dicarboximides used in agriculture. Molecular structures were retrieved from PubChem^123^.

