## Supplementary note 1 for "FungAMR: A comprehensive portrait of antimicrobial resistance mutations in fungi"

### Supplementary note 1–analysis of the AFRbase resource

Recently, Jain, Singhal, and Kumar [1] published a database of fungicide and antifungal resistance genes–AFRbase, obtained *via* textmining. Whilst textmining allows novel inference of biological insight, and has been applied in many fields, AFRbase contains errors.

AFRbase consists of webtables of mutations in genes that confer a resistance against a given antifungal drug. According to the authors, the database consists of both manually curated and computationally asserted records of drug resistance associated mutations. The interpretability, however, of these records is limited, as no minimum inhibitory concentration (MIC) value is reported for a record ([fig:example-record]).

| Features | |
|---|---|
| AFRbase ID | 166 |
| Fungal Pathogen | Aspergillus fumigatus |
| Host | Human |
| Disease caused | NA |
| Mode of transmission | NA |
| GenBank ID | NA |
| Gene ID | 3509526 |
| Gene Name | CYP51a |
| Gene Locus | AFUA_4G06890 |
| Uniprot Entry* | Q4WNT5 |
| Sequence | MVPMLWLTAYMAVAVLTAILLNVVYQLFFRLWNRTEPPMVFHWVPYLGSTISYGIDPYKFFFACREKYGDIFTFILLGQKTTV[illegible] |
| Amino Acid Mutations | P216L |
| Drug against which resistance is gained | Posaconazole |
| Functional Annotation | NA |
| PubMed ID* | 16372009 |
| Reference | 10.3201/eid1507.090043 |

*An example record of AFRbase.*

Furthermore Jain, Singhal, and Kumar [1] does not state which textmining method is employed, and it appears no attempt to analyse the possible errors in AFRbase as been made. For textmining, a proper analysis of errors is a crucial step, as any machine learning method has an error rate. By analysing their database, it is clear that there are a lot of unknowns about the textmining technique .

As this database could be of interest for clinicians and routine clinical analysis it is important that the sources which are cited by the database are accurate; the alternative is a lot of time lost looking for the relevant literature. In AFRbase, 40% of records cite a source from PubMed that does not mention the fungal pathogen stated in the AFRbase record. For some pathogens, such as *Candida tropicalis* this error rate is close to 50% ([fig:topic-correctness]).

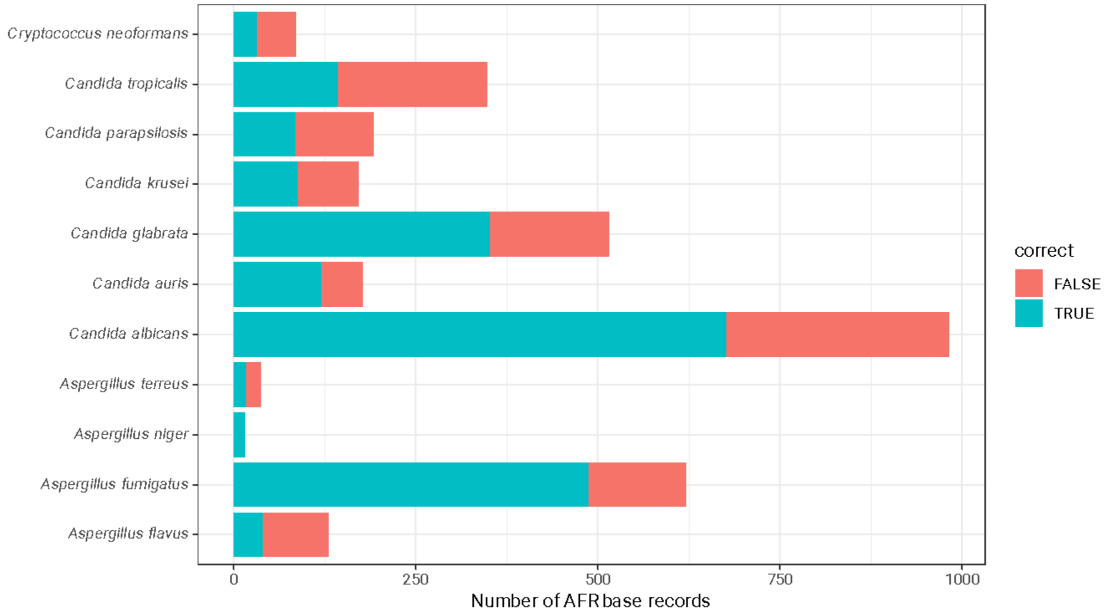


*Number of records in AFRbase which cite a PubMed article that reports on the same organism of the record, split by organism type. If the organism of the record is mentioned in the abstract, it is indicated as 'correct'.*

This finding shows that the AFRbase database has problems with reliability: since the textmining system of AFRbase found words that are not in the pubmed abstracts, it is difficult to interpret the correctness of an AFRbase record.

#### Methods

Using rvest (1.0.4, ) in R (4.4.0, ) the records of the AFRbase were downloaded and converted to YAML files on the 8th of March. Each YAML file contained the records of an AFRbase entry, such as the pmid it cites, and the protein it describes.

These YAML files were loaded into a graph database using rdflib () and pyyaml ( in python. The pubmed identifiers of each AFRbase record were cross-referenced with the pubmed database to download the pubmed data for each article using the metapub () package. This was serialised into a graph structure using rdflib. The final dataset consists of the AFRbase graph information of all the records, and the pubmed graph, that contains the information for each cited pubmed article, such as publication date, abstract text and mesh subject headings.

Using the oxigraph graph database , SPARQL queries were written analyse the correctness of the AFRbase data. The correspondence between the AFRbase 'fungal pathogen' field and the pathogens described in the cited abstract was investigated by searching for the second part of the species name in the xml version of the abstract. For example, if a record on *Candida auris* would be checked, the lower case version of abstract that record cites would be searched for the word 'auris'.

The whole analysis is available as a Snakemake pipeline (5.10.0, ).

`https://github.com/Luke-ebbis/afrbase-analysis` is a snakemake workflow to reproduce the analysis outlined in this information supplement.
